# PLA-complexity of *k*-mer multisets

**DOI:** 10.1101/2024.02.08.579510

**Authors:** Md. Hasin Abrar, Paul Medvedev

## Abstract

**Motivation:** Understanding structural properties of *k*-mer multisets is crucial to designing space-efficient indices to query them. A potentially novel source of structure can be found in the rank function of a *k*-mer multiset. In particular, the rank function of a *k*-mer multiset can be approximated by a piece-wise linear function with very few segments. Such an approximation was shown to speed up suffix array queries and sequence alignment. However, a more comprehensive study of the structure of rank functions of *k*-mer multisets and their potential applications is lacking.

**Results:** We study a measure of a *k*-mer multiset complexity, which we call the PLA-complexity. The PLA-complexity is the number of segments necessary to approximate the rank function of a *k*-mer multiset with a piece-wise linear function so that the maximum error is bounded by a predefined threshold. We describe, implement, and evaluate the PLA-index, which is able to construct, compact, and query a piece-wise linear approximation of the *k*-mer rank function. We examine the PLA-complexity of more than 500 genome spectra and several other genomic multisets. Finally, we show how the PLA-index can be applied to several downstream applications to improve on existing methods: speeding up suffix array queries, decreasing the index memory of a short-read aligner, and decreasing the space of a direct access table of *k*-mer ranks.

**Availability:** The software and reproducibility information is freely available at https://github.com/medvedevgroup/pla-index

## 1 Introduction

Understanding structural properties of *k*-mer multisets is crucial to designing space-efficient indices to query them. Without exploiting such properties, indices cannot hope to overcome worst- or average-case space lower bounds [19]. However, indices can outperform the lower bounds when they exploit the non-uniform structure of the data, e.g. either by parameterizing the theoretical analysis or simply by showing superior performance in practice [17]. For example, the fact that many *k*-mer sets have the spectrum-like property [4] has recently been exploited to design a minimal perfect hash function which substantially outperforms the theoretical lower bound in practice [24].

A potentially novel source of structure can be found in the rank curve of a *k*-mer multiset. The *k-spectrum* of a genome is the multiset of all *k*-mers occurring in it. Let 𝕊 be an array of the lexicographically sorted *k*-spectrum. We think of a *k*-mer as an integer between 0 and 4^*k*^ − 1, encoded in a way so that the numerical order of the integers corresponds to the lexicographical order of the strings. The rank function takes a *k*-mer *x* and returns max_*i*_ 𝕊 [*i*] *< x*. For *x* ∈ 𝕊, this is the first location of *x* in S. Figure 1A) shows a real-world *k*-spectrum– observe how linear-like it appears.

**Fig. 1.**
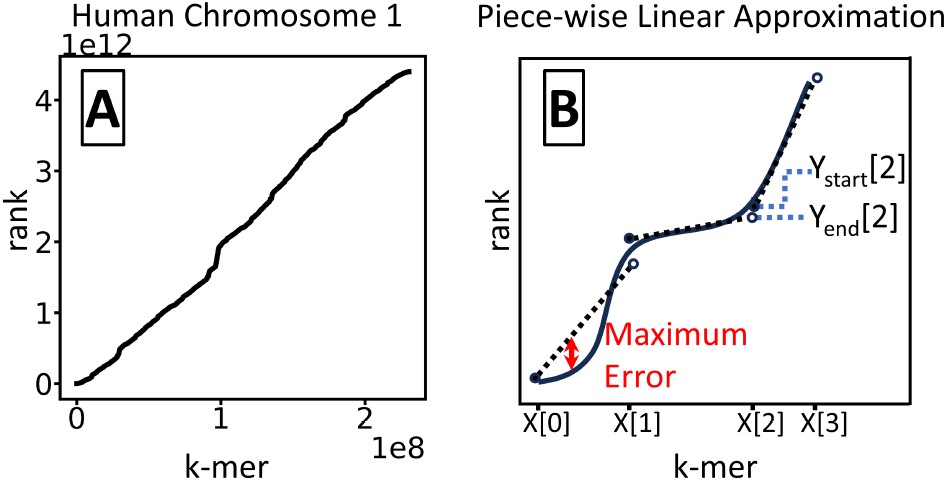
Panel A shows the rank curve for the 21-spectrum of human chr1. Panel B shows a cartoon illustration of a rank curve (solid black curve) and a piece-wise linear approximation (dashed black lines).

A recent line of work applies machine learning to identify and exploit internal patterns in the rank curve of a dataset [16]. Machine learning is used to train a model to fit the curve in a way that minimizes the distance between a *k*-mer’s actual and predicted rank. Such “learned indices” have been designed for genomic datasets [15, 12, 14, 13] and were used to speed-up suffix array search [15] and sequence alignment [14, 13]. However, it was also observed that they do not outperform classical string indices such as tries [7]. Moreover, even though machine learning is able to make use of the structure of *k*-mer spectra, it does not give clear insights into what that structure is.

A simpler and more promising approach is to approximate rank with a piece-wise linear function [10] (see Figure 1B). It was shown that a surprisingly few number of segments are sufficient to obtain a good approximation on real-world data [10, 9, 15]. Moreover, there is an efficient algorithm to find a piece-wise linear approximation (PLA) with a minimum number of segments, for a predefined maximum error threshold [21]. It was therefore suggested by [1] that the number of such segments can serve as a measure of compressibility. We follow this idea and call this the *PLA-complexity* of a *k*-mer multiset. The PLA-complexity is likely connected to the size of the models in the learned index approaches, but we find it a more natural measure of complexity because its definition is simple and does not rely on the output of a black-box machine learning algorithm.

In this paper, we pursue the following questions: (1) what is the PLA-complexity of *k*-mer multisets in practice, (2) can we design compact and efficient *k*-mer indices based on a PLA of a *k*-mer multiset, and (3) can low PLA-complexity be exploited in downstream bioinformatic applications? Some partial answers to these questions are already provided by the Sapling [15] and PGM-index [9] papers. Sapling constructs a data-agnostic PLA that has evenly distributed segments of fixed width. As such, there is no guarantee on the maximum error of the PLA. Nevertheless, they show how it can significantly accelerate suffix array lookups and sequence alignment. They also look at the relationship between the PLA’s error and number of segments, for human chromosome 1. The PGM-index is a more general PLA-based index that computes an optimal segmentation and stores it compactly. Moreover, it uses a recursive approach to represent the starting *k*-mers of the segments themselves using a PLA. Though the PGM-index was not evaluated on *k*-mer multisets, we find it performs well in our experiments and its ideas translate well to the *k*-mer setting.

Here, we undertake a more comprehensive study of the PLA-complexity of *k*-mer multisets. We describe, implement, and evaluate the *PLA-index*, which is able to construct, compact, and query a PLA of the *k*-mer rank function. Our index is similar to the PGM-index, but modifies the construction and compaction method to reduce space. We then examine the PLA-complexity of over 500 genome spectra and several other genomic multisets. Finally, we show how the PLA-index can be applied to several downstream applications to improve on existing methods: speeding up suffix array queries, decreasing the index memory of a short-read aligner, and decreasing the space of a direct access table of *k*-mer ranks.

## 2 Preliminaries

### Multisets of *k*-mers

Let *S* denote a multiset of *k*-mers and let *N* denote the size of *S*. Let *n* denote the number of distinct *k*-mers in *S*. Let 𝕊 denote an array of *N* elements (indexed from 0) which are the sorted elements of the multiset *S*. For example, if we take the multiset of all 2-mers in the string GCCACC, then *S* = {GC, CC, CA, AC, CC}, 𝕊 = (AC, CA, CC, CC, GC), *N* = 5, and *n* = 4.

We distinguish between two ways that S may be represented, which affects the cache locality of accessing its elements. In the *indirect* case, 𝕊is represented indirectly via pointers. For example, let *S* be the spectrum of a genome. Then the suffix array of the genome is an indirect representation of 𝕊 (ignoring suffixes shorter than *k*). That is, accessing the suffix array at location *i* gives you a location in the genome that is the start of the *k*-mer 𝕊 [*i*]. In such a situation, each consecutive access to a *k*-mer of 𝕊 incurs a cache miss. In the *direct* case, 𝕊 is represented directly, e.g. an array of *k*-mers. In such a case, accessing consecutive values of 𝕊 is fast due to cache locality.

For a *k*-mer *x* ∈ *S*, we define rank(*S, x*) as the largest integer 0 ≤ *i < N* such that, for all 0 ≤ *j < i*, 𝕊 [*j*] is strictly less than *x*. For *x* ∈*/ S*, we define rank(*S, x*) = −1. Using our previous example of *S*, rank(*S*, CC) = 2, rank(*S*, AC) = 0, and rank(*S*, AG) = −1.

### Piece-wise linear function

A *piece-wise linear function* is defined by an array of *k*-mers *X*, sorted in increasing order, and two arrays *Y*_start_ and *Y*_end_ of y-values. Intuitively, *X* array represents the breakpoints of the function, *Y*_start_ represents the y-value of the segment starting at a breakpoint, and *Y*_end_ represents the y-value of the segment ending at a breakpoint (Figure 1B). We let *b* denote the number of segments. Formally, for 0 ≤ *i < b*, the *i*^th^ line segment connects the points (*X*[*i*], *Y*_start_[*i*]) and (*X*[*i* + 1], *Y*_end_[*i* + 1]). It has slope 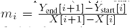. To evaluate the function at a *k*-mer *x*, we first find the largest integer *i* such that *X*[*i*] ≤ *x* and then evaluate *x* using linear interpolation along the *i*^th^ segment. Formally,

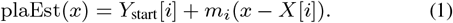

### PLA-complexity

We say that a piece-wise linear function approximates the rank function with a maximum error of *ε* iff for any *k*-mer *x* ∈ *S*, | rank(*S, x*) − plaEst(*x*)| ≤ *ε*. For a given *S*, the *PLA-complexity* of *S* is a function mapping *ε* to the smallest number of segments achievable by a piece-wise linear function with a max error of *ε*. Following the work of [9, 8], we will apply the following model to quantify the PLA-complexity of real-world datasets and help interpret the space and time complexity of our index.

#### Definition 1

(Scaling Model). *Under the Scaling Model, the minimum number of segments b needed to approximate a k-mer multiset by a piece-wise linear function with a maximum error of ε is b* = *Nβ/ε*^*α*^, *where α and β are dataset-specific constants independent of N* . *Moreover, we assume that N/n is a dataset-specific constant independent of n or N* .

Under the Scaling Model, the PLA-complexity of a dataset can be represented by two values, *α* and *β*. We are unaware of any theoretical basis for this model, but it has been supported by experimental data [9, 8].

### Operations

Given a *k*-mer multiset *S* and a *k*-mer *x*, our indices will support two operations:

- SEARCH(*x*) returns a value *i* such that S[*i*] = *x* if *x* ∈ *S* and *i* = −1 otherwise, and
- RANK(*x*) returns rank(*S, x*). ^1^

We stress that our indices are not dictionaries, i.e. they are data structures stored on top of some pre-existing representation of S (for a PLA-based dictionary solution, see [1]).

Indices to support these operations involve an inherent space/time trade-off. Our PLA-index and its variations will fall in between the following two extremes. On one extreme, binary search does need space for an index but is considered slow because it does not have cache locality. At the other extreme, one can construct a minimal perfect hash function (MPHF), with the set of keys being the distinct *k*-mers in *S*. An MPHF is a data structure that maps elements of the key set of size *n* to an integer between 0 and *n* − 1, without any collisions. For *k*-mers that are not in the key set, the MPHF maps them to an arbitrary integer between 0 and *n* − 1. With an MPHF, we can construct a direct-access table (i.e. an array) of size *n* to store the exact rank of each *k*-mer. Such a solution gives very fast queries but uses Θ(*n* lg *N*) space.

### O’Rourke’s algorithm

To compute the piece-wise linear approximation of a rank curve, we make use of an algorithm published by O’Rourke [21] and implemented in [1]. The input to the algorithm is presented in an online manner, i.e. one element at a time. Each element consists of an x-value *x* and a y-range [*𝓁, h*]. It is required that *x* is strictly larger than previous x-values. A line is said to fit these points if for each *x*, its y-value lies in the range [*𝓁, h*]. The algorithm maintains the set of all lines that fit the input so far^2^. When an input element is presented such that there is no longer any line that fits the data, the algorithm outputs a line that fits the previous ranges and terminates. The algorithm running time and memory is linear in the number of input elements.

## 3 The basic PLA-index

The basic PLA-index is constructed from a sorted array S of *N k*-mers and an error threshold *ε*. It consists of a piece-wise linear function, stored in *X, Y*_start_, and *Y*_end_ arrays, and a *shortcut array D*.

### Construction

We process the *k*-mers of S from smallest to largest, treating each *k*-mer as an integer between 0 and 4^*k*^ − 1. For each *k*-mer *x* ∈ 𝕊, we define its *y*-range to be [rank(*S, x*) − *ε*, rank(*S, x*) + *ε*]. We feed *x* and its *y*-range to O’Rourke’s algorithm, until the algorithm stops and outputs a line that fits the previously given ranges. We store the *k*-mer and the y-value of the first point of this line in *X* and *Y*_start_, respectively. We store the y-value of the last fitted point in *Y*_end_. We then restart O’Rourke’s algorithm, but we reuse the last *k*-mer of the previous fitted line to start the next iteration. As will be clear later, this will allow us to more compactly represent the *Y*_end_ array. The construction algorithm runs in Θ(*N*) time and uses memory linear in the maximum number of *k*-mers in a fitted line.

In order to store the *Y*_start_ and *Y*_end_ values as integers, rather than reals, we round the y-values returned by O’Rourke’s algorithm. In some rare cases, this results in a line that no longer fits the range of some point *x*. In these cases, we re-run O’Rourke’s algorithm but manually terminate it at *x*, using whatever line is a valid fit up to that point. We then restart from *x*. We find in practice that the amount of such forced breaks is negligible.

We also construct a shortcut array *D*, which will be used to speed up binary search on *X. D* contains 2^*𝓁*^ elements, where *𝓁 <* 2*k* is a parameter of the algorithm. In practice, we set *𝓁* = 16. The *i*^th^ element of *D* is the smallest position *j* such that the first *𝓁* bits of *X*[*j*] are at least *i*.

### SEARCH and RANK queries

Let *x*_*𝓁*_ be the first *𝓁* bits of *x*, viewed as an integer. We do a binary search in *X* between positions *D*[*x*_*𝓁*_] and *D*[*x*_*𝓁*_ + 1] to find the largest index *i* such that *X*[*i*] ≤ *x*. Then we can compute plaEst(*x*) according to Eq. 1. We then do a binary search in 𝕊 [⌊plaEst(*x*) − *ε*⌋], …, 𝕊 [⌈plaEst(*x*) + *ε*⌉] to find the smallest value *i* such that 𝕊 [*i*] ≥ *x*. We return *i* if 𝕊 [*i*] = *x* and return -1 otherwise. The only difference between SEARCH and RANK is that with SEARCH we can shortcut the binary search in S as soon as we hit a value *i* such that 𝕊 [*i*] = *x*.

### Time complexity

The worst-case time complexity of computing plaEst is Θ(lg *b*) and extending it to the full RANK operation adds another Θ(lg *ε*) time, where *b* is the number of segments. Thus the worst case total time for RANK is Θ(lg *b* + lg *ε*). Under the Scaling Model, the time becomes Θ(lg *N* − (*α* − 1) lg *ε*). This reduces the runtime compared to that of the Θ(lg *N*) index-less binary search (note that a result of [8] states that *α* ≥ 1 for all inputs).

However, the big advantage is gained from cache effects, since in practice we observe that the PLA-index fits into the cache while 𝕊 is stored in RAM. In this case, computing plaEst(*x*) only uses cache accesses and takes Θ(lg *b*) = Θ(lg *N* − *α* lg *ε*) time, while the remaining binary search of RANK, which accesses RAM, takes Θ(lg *ε*) time. In other words, the number of RAM accesses now depends only on *ε* and not on *N* .

### Compact storage

The arrays *X, Y*_start_, *Y*_end_, and *D* can be naïvely stored using 2*kb, b* lg *N, b* lg *N*, and 2^*𝓁*^ lg *b* bits, respectively. However, we exploit properties of the first three arrays to store them more compactly. The *X* array is an array of increasing integers, which can be represented compactly using the Elias-Fano technique [5, 6], as described in [26] and implemented in [22]. Elias-Fano encodes an array of *m* non-decreasing elements coming from a universe of size *U* in *m*⌈lg(*U/m*) + *c*_ef_⌉ + *o*(*m*) bits, where *c*_ef_ is a number between 1.5 and 2. It supports constant time access to arbitrary elements. In our case, the space for the X-values is *b*⌈lg(4^*k*^*/b*) + *c*_ef_⌉ + *o*(*b*) bits.

Elias-Fano lookups are nevertheless slower than the naïve encoding in practice, and we query *X* repeatedly as part of computing plaEst. We therefore consider an alternate encoding in practice. Consider the *i*^th^ element and let *x* = *X*[*i*]. If the values of *X* were distributed evenly among all the universe of 4^*k*^ *k*-mers, then *x* would be *i*4^*k*^*/b*. Instead of storing *x*, we store the difference between *x* and this value, i.e. we store *x* − *i*4^*k*^*/b*. To the extent that the *k*-mers of *X* are somewhat evenly distributed among the universe, the stored difference is small. This allows us to use a small number of bits to store each value. We use a constant-width encoding, where the number of bits is chosen so it can fit the largest difference in *X*. We find in practice that this takes more space than Elias-Fano but less space than the naïve 2*k* bit encoding, while performing lookups as fast as with the naïve encoding.

To compact the *Y*_start_ array, we first show that its values are non-decreasing. The following lemma gives the basis for this.

#### Lemma 1.

*Given a line fitted by O’Rourke’s algorithm, let y*_start_ *denote the y-value of the line at the first fitted point and let y*_end_ *denote the y-value of the line at the last fitted point. Let X*_fit_ = (*x*_start_, …, *x*_end_) *be the sequence of k-mers that are covered by a run of O’Rourke’s algorithm during the PLA-index construction algorithm. Then there exists a line that fits X*_fit_ *such that*

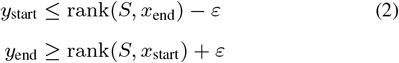

Proof. Observe that if rank(*S, x*_start_) + *ε* ≤ rank(*S, x*_end_) − *ε* then any fitted line will satisfy the lemma. Therefore, we assume that rank(*S, x*_end_) *<* rank(*S, x*_start_) + 2*ε*. Consider the line with *y*_start_ = rank(*S, x*_end_) − *ε* and *y*_end_ = rank(*S, x*_start_) + *ε*. Let *x* be an element of (*x*_start_, …, *x*_end_) and let *y* be the value of this line at *x*. By our assumption, the line has positive slope, and, therefore, *y*_start_ ≤ *y* ≤ *y*_end_. Then,

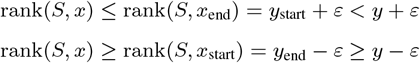

Therefore, the line covers *x* and hence fits *X*_fit_. Figure 2 illustrates the idea of the proof.

**Fig. 2.**
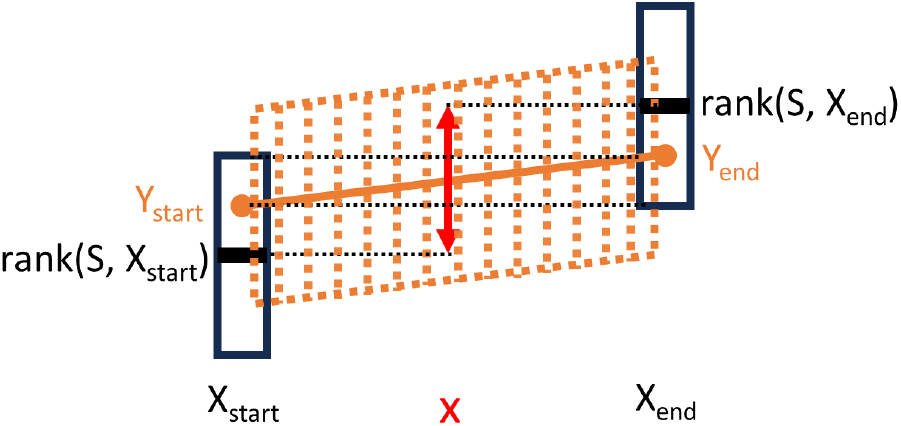
Illustration of the proof of Lemma 1. The values of rank(*S, x*_start_) and rank(*S, x*_end_) are shown in black, along with the *±ε* vertical region around them. The proposed fitting line is shown in solid orange, and the dashed region denotes the *±ε* region covered by the line. A sample middle point *x* is shown in red and its possible rank(*S, x*) values are denoted by the range of the red arrow. The proof is based on the fact that the red range is covered by the orange range.

Lemma 1 can be used to guarantee that both *Y*_start_ and *Y*_end_ are non-decreasing, but our encoding only needs that *Y*_start_ is non-decreasing. To guarantee that the line chosen by O’Rourke’s algorithm satisfies the *y*_start_ constraint (Eq. 2), we can choose the line with the smallest *y*_start_ value. We now prove that *Y*_start_ is non-decreasing:

#### Corollary 1.

*Let* 0 ≤ *i < b* − 1. *Then Y*_*start*_[*i*] ≤ *Y*_*start*_[*i* + 1].

Proof. Let *x* be the last *k*-mer fitted by segment *i* (i.e. the segment starting at *X*[*i*]). Because *x* is the end of segment *i*, Lemma 1 gives that *Y*_start_[*i*] ≤ rank(*S, x*) − *ε*. In the way that we use O’Rourke’s algorithm during construction, we have that *x* = *X*[*i* + 1]. However, all that is needed for the proof is that *x* ≤ *X*[*i* + 1] and, since the rank function is increasing, rank(*S, x*) ≤ rank(*S, X*[*i* + 1]). Therefore, *Y*_start_[*i*] ≤ rank(*S, X*[*i* +1]) − *ε*. Simultaneously, since segment *i* +1 must fit *X*[*i* + 1], we have that *Y*_start_[*i* + 1] ≥ rank(*S, X*[*i* + 1]) − *ε*. Hence, we get *Y*_start_[*i*] ≤ rank(*S, X*[*i* + 1]) − *ε* ≤ *Y*_start_[*i* + 1].

Therefore, we can encode the *Y*_start_ values using Elias-Fano. *Y*_start_ contains *b* elements from a universe of size *N*, so the space used is *b*⌈lg(*N/b*) + *c*_ef_⌉ + *o*(*b*) bits. Note also that the *Y*_start_ values are only accessed once during a rank operation, to compute the slope of the segment. Therefore, the slower access time of Elias-Fano does not have a substantial affect on runtime.

To compact the *Y*_end_ array, we encode each value relative to *Y*_start_. Consider be an arbitrary position *i* in *Y*_end_ and let *x* = *X*[*i*] be the *k*-mer at *i*. Recall that the segment starting at *X*[*i* − 1] ends at *x* with a y-value of *Y*_end_[*i*], while the segment starting at *x* starts with a y-value of *Y*_start_[*i*]. Both segments are subject to the max error constraint at *x* so therefore |*Y*_end_[*i*] − rank(*S, x*)| ≤ *ε* and |*Y*_start_[*i*] − rank(*S, x*)| ≤ *ε*. Putting it together, −2*ε* ≤ *Y*_end_[*i*] − *Y*_start_[*i*] ≤ 2*ε*. Therefore, we can encode *Y*_end_[*i*] by encoding the difference with *Y*_start_[*i*], using only ⌈lg(1 + 4*ε*)⌉ bits per entry. Note that this is the reason why we run O’Rourke’s algorithm starting from the previous fitted *k*-mer, so that we can guarantee that we can encode the difference between *Y*_start_ and *Y*_end_ efficiently using a fixed number of bits. Finally, we can state the space usage of PLA-index:

#### Theorem 1.

*Assuming the use of Elias-Fano encoding for the X array, the total bits used by the basic PLA-index is*

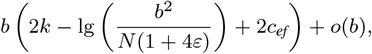

*where c*_*ef*_ *is a value between 1*.*5 and 2. If we assume the Scaling Model and that the number of segments in the PLA-index is the same as in a PLA with a minimum number of segments, then the bits used is*

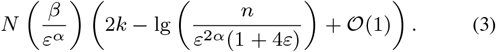

We observe that in Eq. 3, the bits used per segment decrease as *n* increases, though the total space still increases with *n*.

### Choosing *ε*

If the user has in mind how large of an error their runtime can tolerate, they can set *ε* directly. Alternatively, they can set a target memory for the index. In this case, we can run the construction algorithm and measure the index size with *ε* = 1. Using the scaling model, we can solve for *β* and then use Theorem 1 with *α* = 1 to extrapolate what *ε* should be in order to achieve the target memory. This approach is not very precise but can get the index in the ballpark of the target memory. More sophisticated techniques have been presented in [10], but we have not implemented them in our prototype.

## 4 PLA-index with repeat stretching

In this section, we describe an alternative version of the PLA-index which is preferable when the common query is SEARCH rather than RANK. The idea is that if we are allowed to report any position containing *x*, rather than necessarily the first one, then we can allow plaEst(*x*) to be *ε* higher than the last (rather than the first) occurrence of *x*. In this way, we can give more leeway to O’Rourke’s algorithm, allowing it to use fewer segments.

### Construction and storage

Let occ(*x*) define the number of times a *k*-mer is repeated in S. We modify the basic PLA-index construction algorithm by modifying the y-ranges fed to O’Rourke’s algorithm, making them [rank(*S, x*) − *ε*, rank(*S, x*) + *ε* + occ(*x*) − 1]. In other words, we expand the y-range so that any y-value in the range is at most *ε* away from some position of *x*, but not necessarily the first one.

When using repeat stretching, we can no longer guarantee that *Y*_end_[*i*] and *Y*_start_[*i*] values lie within 2*ε* of each other, because occ can be as high as *N* − *n* + 1. Nevertheless, we observe that |*Y*_end_[*i*] − *Y*_start_[*i*]| still tends to be small. We therefore can encode *Y*_end_ using a technique for variable-width encoding of integers that allows random access, known as Directly Addressable Codes [3] and implemented in [11]. Since the encoding only works for non-negative values, we transform the values prior to encoding to be 2|*Y*_end_[*i*] − *Y*_start_[*i*]| + *t*, where *t* = 1 if *Y*_end_[*i*] − *Y*_start_[*i*] is positive and *t* = 0 otherwise.

The reason that the basic PLA-index construction algorithm reused the previously fitted *k*-mer for O’Rourke’s algorithm was to guarantee that *Y*_end_[*i*] − *Y*_start_[*i*] is bounded. Since this is no longer possible, we now do not reuse the *k*-mer, thereby decreasing the number of segments. Further, the proof of Corollary 1 also works without the *k*-mer reuse, and so we can encode *Y*_start_ as before.

### SEARCH and RANK queries

The algorithm for the SEARCH query remains the same as for the basic PLA-index. Let *p* = SEARCH(*x*). For the RANK query, we further need to continuously decrement *p* as long as 𝕊 [*p*] = *x*.

If 𝕊 is stored directly, then RANK is fast because of cache locality. If 𝕊 is stored indirectly, then each access to a *k*-mer of 𝕊 incurs a cache miss, making RANK slow. This can be sped by storing an additional bitvector *B* which marks the positions in S which have a *k*-mer that is different from the preceding *k*-mer. However, this adds *N* bits of space, which can easily dwarf the space of the PLA-index. We therefore do not recommend using repeat stretching for the case that RANK needs to be supported and S is stored indirectly.

## 5 PLA-index-exact

The basic (and repeat-stretched) PLA-index can be viewed as using a tiny bit more space to significantly speed up binary search. However, it still does not perform as fast as a direct access table of *k*-mer ranks. We therefore propose a variant of the PLA-index that is nearly as fast as the direct access table of *k*-mer ranks but takes substantially less space. A similar idea was used in the context of a PLA-based dictionary [1].

As with the direct access table of *k*-mer ranks solution, we construct an MPHF with the set of keys being the distinct *k*-mers in *S*. We then build the basic PLA-index. Finally, we construct an error array *E* of size *n* that, for each *x* ∈ *S*, sets *E*[MPHF(*x*)] to be plaEst(*x*) − rank(*S, x*). To perform RANK(*x*), we let *p* = plaEst(*x*) − *E*[MPHF(*x*)], check if 𝕊 [*p*] = *x*, and if yes, then return *p*, otherwise return -1. The binary search through S done by the basic PLA-index is now avoided and replaced with the cost of one MPHF calculation and one access to *E*.

Since each entry in *E* is guaranteed to be between −*ε* and +*ε*, the additional space required over the basic PLA-index is *n* lg(2*ε* + 1) bits for the *E* array and the space to store the MPHF (usually 2-4 bits per *k*-mer). The additional space is decreased for lower *ε* but the space of the basic PLA-index is increased (Theorem 1). We explore the trade-off in Section 6.4.

## 6 Experimental results

In this section, we demonstrate how PLA-complexity can be exploited for a variety of bioinformatic applications through the PLA-index. We also empirically study the PLA-complexity of *k*-mer and related multisets. PLA-index is available free and open-source at https://github.com/medvedevgroup/pla-index, along with reproducibility information for the experiment in this section.

### 6.1 Experimental setup

We used a machine with an Intel(R) Xeon(R) CPU E5-2683 v4 @ 2.10GHz processor with 64 cores and 512 GB of memory to run our experiments. Unless otherwise stated, all reported running times are medians of five runs. The results of all SEARCH and RANK operations were confirmed for correctness. We used *k* = 21 for all indices, in line with previous work [15]. We used libdivsufsort [18] to construct suffix trees and we use PTHash [25] for MPHF construction.

### 6.2 PLA-index speeds up suffix array queries

We demonstrate how the repeat-stretched PLA-index can be used to speed up the SEARCH query in the case that S is represented indirectly via a suffix array of the hg38 human genome. We compare against an index-less binary search, Sapling [15], and PGM-index [9] (Table 1). First, we observe that PLA-index nearly halves the time of a regular binary search when used with *ε* = 15, for only 50MiB of space. Second, when compared to Sapling, for similar index size, PLA-index is at least 24% faster (a precise comparison is difficult because the index sizes do not match exactly). These results indicate the importance that an error guarantee, optimal segment selection, and compact representation has for rank estimation, as these are the main improvements of PLA-index over Sapling. Third, PLA-index is slightly smaller but slightly slower then the PGM-index. Finally, we see that reducing *ε* starts to have negligible runtime benefits beyond *ε* = 63, likely due to binary search no longer being the bottleneck of the query algorithm.

**Table 1.** Suffix array search on the human whole genome (hg38). We randomly chose 50 million positions of hg38 and used the *k*-mers starting at those positions as queries. We measure total time of all 50 million queries. PGM-index only returns an interval of possible locations, so we supplemented it with the same binary search as we have in PLA-index.

| Method | Max error | N. segments | Index Construction Time (s) | Index Size (MiB) | SEARCH Time (s) |
| --- | --- | --- | --- | --- | --- |
| Repeat-stretched PLA-index | 1,023 | 269,717 | 1,105 | 4 | 108 |
|  | 255 | 1,066,934 | 1,106 | 13 | 93 |
|  | 63 | 4,290,786 | 1,085 | 50 | 85 |
|  | 15 | 18,497,919 | 1,139 | 213 | 82 |
| PGM-index | 1,023 | N/A | 1,064 | 5 | 106 |
|  | 255 | N/A | 1,083 | 21 | 89 |
|  | 63 | N/A | 1,095 | 85 | 73 |
|  | 15 | N/A | 1,119 | 370 | 60 |
| Sapling | N/A | 1,048,576 | 1,869 | 16 | 121 |
|  |  | 2,097,152 | 1,842 | 32 | 117 |
|  |  | 4,194,304 | 1,846 | 64 | 112 |
|  |  | 8,388,608 | 1,966 | 128 | 120 |
| Binary Search | N/A | N/A | N/A | 0 | 153 |

We find it remarkable how small the PLA-index is. With only 4 MiB of memory, we can estimate the position of a *k*-mer in a suffix array of size ≈ 3 billion to within 1023 positions. With 50 MiB of memory, we can estimate it to within 63 positions.

### 6.3 PLA-index reduces memory use of read aligner

Strobealign is a recent aligner for short reads [31]. To represent the reference, it uses a seed table where each row corresponds to a seed and contains the 64-bit hash of the seed sequence and its associated data. The rows are sorted in increasing order of hash values. Thus, the seed table is a direct representation of a multiset *S*, with the minor difference that each element is not a *k*-mer but a 64-bit hash value. To align the reads, strobealign repeatedly searches the table for a specific seed hash value. In order to avoid cache-unfriendly binary search, strobealign also stores a large pointer vector (e.g. for the human reference, it has 2^28^ elements), where the element at position *h* is the index of the first row in the seed table whose hash value starts with *h*. We modified strobealign by replacing the pointer vector with the repeat-stretched PLA-index.

Table 2 shows that the PLA-index takes two or three orders of magnitude less space than the pointer vector. For example, while the pointer vector takes 2 GiB for the human, PLA-index takes 1 MiB for *ε* = 63. The overall memory usage of strobealign is still dominated by other components of their index; though these can be substantially optimized [32], it is outside the scope of our project. We did observe a 1% slow down on Drosophila and about a 5% slow down on human. We believe that this is simply due to the fact that strobealign code has been highly optimized for speed [32], while our implementation is only a prototype. We note that increasing *ε* does not increase the runtime, indicating that the overhead of implementing RANK on a repeat-stretched PLA-index when S is stored directly is negligible.

**Table 2.** Breakdown of memory usage and runtime by Strobealign, with and without our PLA-index. Our modified version replaces the pointer vector with PLA-index. The “Other” column refers to the sum of all other components of the index, present before and after our modifications. To measure runtime, we used 16 threads and measure wall clock time of a single run. The Drosophila experiment uses a read set with 10.5 million paired-end reads and BDGP6.22 [29] as the reference. The human experiment uses a read set with 10 million paired-end reads with T2T-CHM13v2.0 [30] as the reference. All reads were simulated using the same setup as [31].

| Dataset | Max Error | N. segments | Index Memory (MiB) |  |  | RANK runtime (s) |  |
| --- | --- | --- | --- | --- | --- | --- | --- |
|  |  |  | PLA-index | Pointer vector | Other | Modified | Original |
| Drosophila | 255 | 495 | .01 |  |  | 424 |  |
|  | 63 | 6,368 | .08 | 64 | 615 | 420 | 417 |
|  | 15 | 62,092 | .64 |  |  | 420 |  |
| Human | 255 | 28,374 | .3 |  |  | 633 |  |
|  | 63 | 136,322 | 1.5 | 2,048 | 12,710 | 615 | 590 |
|  | 15 | 1,055,142 | 10.6 |  |  | 625 |  |

### 6.4 PLA-index-exact reduces memory of a direct access rank table

We compare PLA-index-exact to the fastest alternative option, which uses the same MPHF as PLA-index-exact but in the table stores each *k*-mer’s exact rank, rather than its error. Table 3 shows that PLA-index-exact offers a drastic improvement, reducing the total space by 61% while keeping the running time virtually the same (using *ε* = 1023 on hg38). PLA-index-exact also more than halves the look up time of the repeat-stretched PLA-index (Table 1), though at a considerable space cost. Table 3 also shows the speed/memory trade-off regulated by *ε*. The run time seems to be influenced by low-level system effects, and we do not pursue the question further in this paper.

**Table 3.** Performance of PLA-index-exact on hg38. We compare against storing the exact ranks in a table, using the 21-spectrum of hg38. The run-times reported are the medians of five independent runs. The queries are the same as in Table 1.

| Max Error | MPHF (MiB) | PLA-index-exact |  |  | Rank table |  |
| --- | --- | --- | --- | --- | --- | --- |
|  |  | PLA-index (MiB) | Table (MiB) | SEARCH time (s) | Table | SEARCH time (s) |
| 16,383 |  | 0.2 | 4,172 | 29 |  |  |
| 4,095 | 728 | 0.8 | 3,615 | 29 | 8,900 | 32 |
| 1,023 |  | 2.9 | 3,059 | 31 |  |  |
| 255 |  | 11 | 2,502 | 35 |  |  |

### 6.5 Repeat stretching helps

Repeat stretching involves a trade-off between reducing the number of segments and using a less space-efficient encoding of *Y*_end_. Table 4 shows that repeat stretching reduces the overall memory, with an increased benefit for smaller *ε*. For example, for *ε* = 15, the size of the index is reduced by 7%. The percent reduction in the number of segments similarly increases as *ε* decreases, with a reduction of 12% when *ε* = 15. On the other hand, the extra bits needed for each *Y*_end_ entry stays constant at 3 bits, for all the tested values of *ε*.

**Table 4.** Comparison of space usage of the basic PLA-index with the repeat-stretched PLA-index on chr1 of hg38. The *Y*_end_ column shows the average number of bits used per entry. The chr1 21-spectrum has *N* = 230, 481, 012 entries, of which *n* = 194, 598, 515 are distinct.

| Error | Basic PLA-index |  |  | With repeat stretching |  |  |
| --- | --- | --- | --- | --- | --- | --- |
| | N. segments | Mem (Mb) | $Y_{\text{end}}$ (bits) | N. segments | Mem (Mb) | $Y_{\text{end}}$ (bits) |
| 1023 | 20,881 | 0.67 | 12 | 19,491 | 0.66 | 15 |
| 255 | 87,999 | 1.16 | 10 | 80,607 | 1.13 | 13 |
| 63 | 372,284 | 3.1 | 8 | 330,969 | 2.9 | 11 |
| 15 | 1,635,558 | 11.2 | 6 | 1,438,552 | 10.3 | 9 |

### 6.6 Empirical study of PLA-complexity

To explore the PLA-complexity across a diverse set of genomes, we downloaded a sample of RefSeq genomes that are complete and full, do not have any missing bases, and are longer than 10,000nt. Our dataset contained 549 genomes, representing the kingdoms of Virus, Bacteria, Archaea, and Fungi. The median genome length was 2.7 mil and the maximum was 63 mil. We used O’Rourke’s algorithm, with *k* = 21, to obtain the PLA-complexity *b* for *ε* ∈ {1, 2, 4, 8, …, 1024}. We then fit a two parameter curve *b* = *βN/ε*^*α*^ to each genome using non-linear least squares regression (function nls in R). As a seed, we set *α* = 1 and *βN* equal to the number of segments for *ε* = 1. We found the fits to be fairly accurate (see for example Figure 3), albeit generally underpredicting the number of segments when the number is small (i.e. for large *ε*). Further details of the fits are available at our GitHub page.

**Fig. 3.**
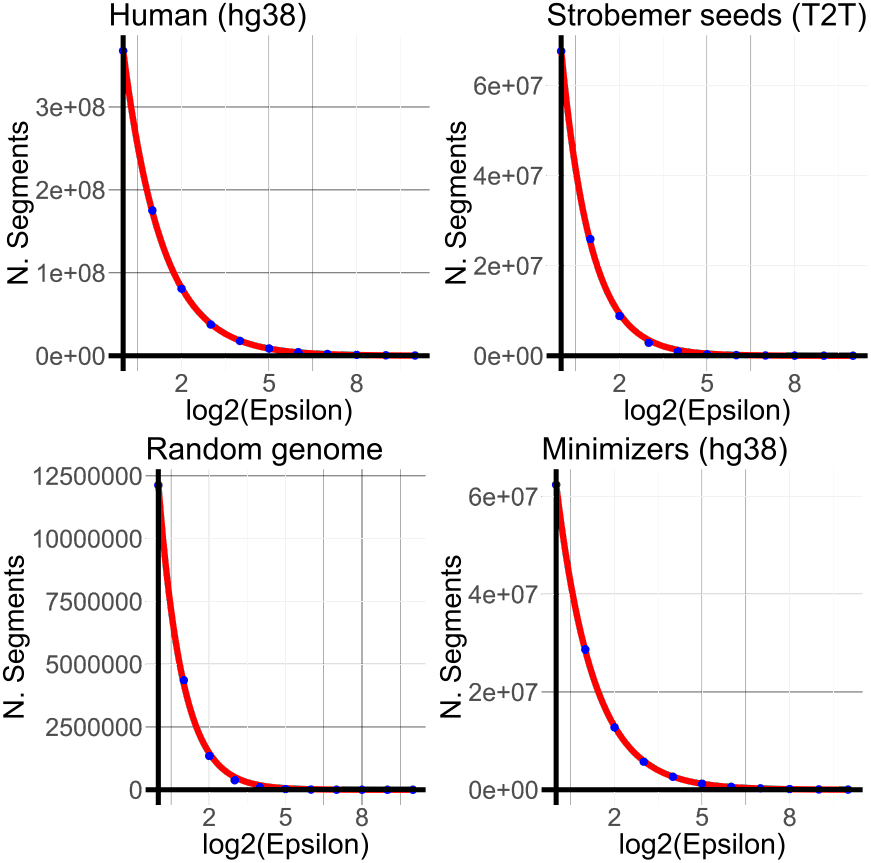
The PLA-complexity curve of some of the datasets from Table 6. The blue dots represent the true number of segments for *ε* = 2^*i*^ for 1 *≤ i ≤* 10. The red line is the best fit of *b* = *Nβ/ε*^*α*^.

We find it striking how tight the distributions of the fitted parameters are (Table 5) . For 95% of the genomes, 1.1 ≤ *α* ≤ 1.2, and, similarly, 0.14 ≤ *β* ≤ 0.17. We did not observe any significant differences between the various kingdoms. We did not find any correlation (i.e. all *R*^2^ *<* 0.02) between either *α* or *β* and the length of the genome, the assembly N50, or the repetitiveness of the genome as measured by *N/n*.

**Table 5.** The fitted PLA-complexity of our RefSeq genome dataset. We show the number of genomes (N.) in each category, the mean and the 5% and 95% percentiles of the fitted values. The first row shows the values for all the genomes combined, while the later rows break them down by Kingdom.

| Category | N. | Fitted degree ($\alpha$) | | | Fitted constant ($\beta$) | | |
| --- | --- | --- | --- | --- | --- | --- | --- |
|  |  | 5th% | Mean | 95th% | 5th% | Mean | 95th% |
| RefSeq dataset | 549 | 1.1 | 1.2 | 1.3 | .14 | .15 | .17 |
| Virus | 127 | 1.1 | 1.2 | 1.3 | .13 | .15 | .17 |
| Bacteria | 200 | 1.1 | 1.2 | 1.3 | .14 | .15 | .17 |
| Archaea | 200 | 1.1 | 1.2 | 1.3 | .14 | .15 | .17 |
| Fungi | 22 | 1.1 | 1.2 | 1.3 | .13 | .14 | .15 |

**Table 6.**
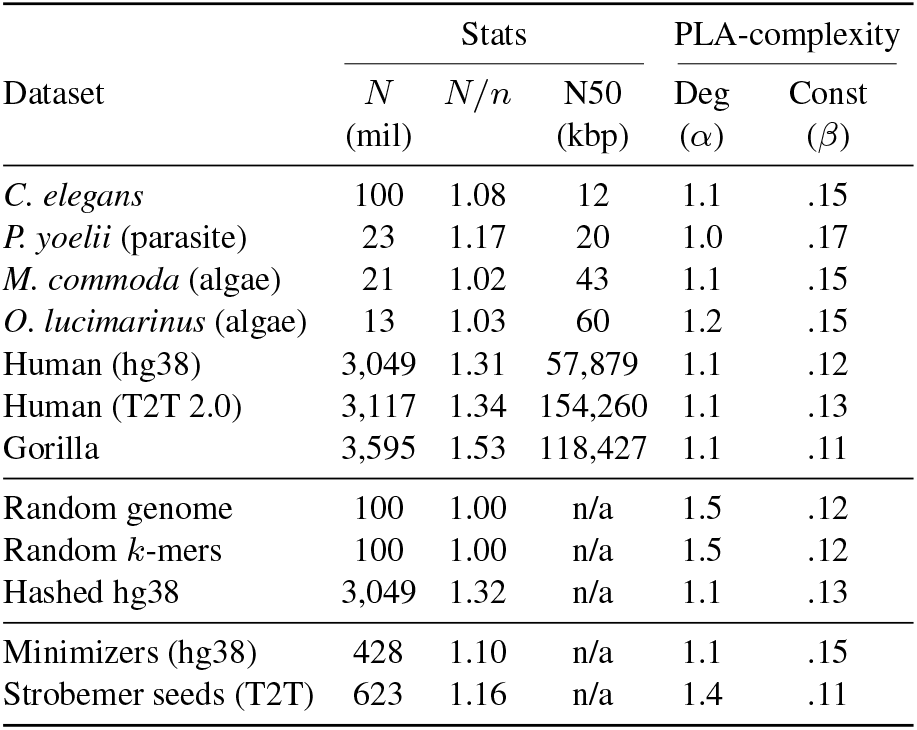
The fitted PLA-complexity of various datasets. The full curves of some of these are visualized in Figure 3. Each of the first 7 rows represents the 21-spectrum from the corresponding genome (accession numbers are available in our GitHub page). The “Random genome” row is the 21-spectrum of a sequence chosen uniformly at random over the DNA alphabet. The “Random *k*-mers” row is a multiset of 21-mers where each 21-mer is chosen uniformly at random from the universe of all 21-mers, with replacement. The “Hashed hg38” row is the multiset of hashed *k*-mers of the spectrum of hg38. The *N* column is the number of elements in the multiset from which the PLA-index is constructed.

Next, we looked at the PLA-complexity of larger genomes (top group in Table 6 and Figure 3). This includes both hg38 and T2T versions of the human, the Gorilla, two plants, a parasite, and *C. elegans*. The fitted degree and constant was mostly within the same range as for the RefSeq genomes. There was one notable outlier, a rodent malaria parasite which had degree *α* = 1.0; however, we were not able to identify anything particular about this assembly or species that could explain the lower *α* value.

Next, we wanted to understand the extent to which the structure of genomic spectra affects their PLA-complexity (Table 6). We found that for a random multiset of *k*-mers, *α* = 1.5. which is substantially higher than the range of 1.0 - 1.3 we found in genomic spectra. There are at least two factors which distinguish a genome spectrum from a random multiset. One is that the *k*-mers of a genome spectra must be overlapping, i.e. not all multisets of *k*-mers can be put together into a continuous genome sequence, even allowing for chromosome breaks. The second is that genomes have repeats, which are very rare in a random multiset of *k*-mers. To tease out the affect of these two, we measured the PLA-complexity of two artificial multisets. For the spectrum of a long random DNA sequence (which has the first but not the second property, since long random strings have a negligible amount of repeats), *α* = 1.5. For the randomly hashed *k*-mers of the hg38 spectrum (which has the second but not the first property, since overlaps between *k*-mers are destroyed after hashing), *α* = 1.1. These results lead us to hypothesize that the salient property of genomic spectra is the presence of repeats, rather than overlapping *k*-mers.

Next, we looked at the PLA-complexity of a non-spectrum *k*-mer multiset. We took the multiset of all minimizers from hg38 (as computed by UST [27] and SSHash [23] using window size of 31 and minimizer size of 21). Note that the lexicographical order in which we store the multiset of minimizers is different from the random ordering used for the minimizer scheme. The fitted degree (*α* = 1.1) is the same as the hg38 spectrum but the constant (*β* = 0.15) is higher then hg38 (*α* = 0.12) (Table 6). It is not clear if this increase is related to the nature of the minimizer sketch or just a fluctuation similar to these we observed in the RefSeq data (Table 5).

Finally, we looked at the PLA-complexity of multisets that are not *k*-mers. We took the multiset of seeds produced by Strobealign [31], run on the T2T genome with default parameters. The fitted degree is *α* = 1.4 (Table 6) is outside the range of our genomic spectra but is similar to a random set of *k*-mers. This is not surprising, as each seed is a random hash of a strobemer sequence.

## 7 Conclusion

In this paper, we initiated the study of PLA-complexity of *k*-mer multisets. We empirically studied the PLA-complexity of genomic *k*-mer spectra as well as other types of genomic multisets, such as a genome’s minimizers or seeds used for alignment. We developed several variations of an index, called the PLA-index, which exploit the small PLA-complexity of genomic data in order to speed up search and rank queries. We demonstrated how the PLA-index can be applied in various settings to achieve dramatic time and/or memory improvements. For example, the PLA-index sped up the binary search for a *k*-mer in a suffix array by two-fold, reduced the memory of a short-read aligner by 2GiB on a human, and reduced the memory of a direct access table of *k*-mer ranks by 61%.

To the best of our knowledge, our paper is the first to study the PLA-complexity across a broad spectrum of genomic datasets. Our main finding is that the PLA-complexities of genome spectra lie in a small range. Beyond this, our study opens more questions than it answers. One of the most basic questions on the theoretical side is to derive the expected number of segments on a random *k*-mer multiset, as a function of *ε*. Some general bounds are already known [8], but they are not informative in our setting. Moving to real genomes, the theoretical questions are likely to become more difficult to answer. It would also be interesting to explore the connection between PLA-complexity and other string complexity measures [20]. In particular, the delta measure [28] seems somehow related and has already been explored for genome strings [2].

Empirically, are our observed fluctuations in the value of the fitted constant *β* just noise, or do they represent some kind of intrinsic property of the *k*-mer multiset input? In contrast, our empirical results do show that the fitted degree *α* captures an important property of genomic spectra that differentiates it from the spectrum of a random sequence. Furthermore, our results only scratch the surface of the PLA-compelxity of non-spectrum datasets, e.g. minimizer multisets or *k*-mer multisets that have been hashed.

One can imagine many ways that low PLA-complexity can be exploited to index *k*-mer multisets. The PLA-index is just one possibility, guided by our own design choices. However, several reasonable alternatives might be pursued. For example, one can abandon the maximum error guarantee of the piece-wise linear function and instead take a heuristic or probabilistic approach to reducing the error (e.g. reduce the average error). While some alternatives were explored in previous works (e.g. [15]), we believe there remains a lot of unexplored potential.

More broadly, low PLA-complexity of genomic spectra could be exploited by other data structures and algorithms. For example, [1] proposed a PLA-based dictionary, but it has not been applied in the *k*-mer setting. Other possibilities include minimum perfect hash functions, rank and select data structures, and *k*-mer counting. We believe that PLA-based approaches have the potential to outperform many state-of-the-art approaches when dealing with genomic *k*-mer data.

## Acknowledgements

We thank K. Sahlin and M. Martin for their help in identifying the Strobealign application and helping guide its implementation. We also thank Giorgio Vinciguerra for helpful discussions.

## Footnotes

1 Note that RANK returns -1 for elements not in *S*. Support for RANK on elements not in *S* could be added to our indices, though we have not implemented it and therefore do not describe the details.

2 The crux of O’Rourke’s algorithm is how to maintain this set efficiently, but since we use the algorithm in a black-box manner, we omit this description.

